# IMMUNE SYSTEM CHALLENGE IMPROVES COGNITIVE-BEHAVIOURAL RESPONSES AND REVERSES MALARIA-INDUCED COGNITIVE IMPAIRMENT IN MICE

**DOI:** 10.1101/2019.12.13.874420

**Authors:** Luciana Pereira de Sousa Vieira, Flávia Lima Ribeiro-Gomes, Roberto Farina de Almeida, Tadeu Mello e Souza, Guilherme Loureiro Werneck, Diogo Onofre Gomes de Souza, Cláudio Tadeu Daniel-Ribeiro

**Author notes:** Corresponding author at *Laboratório de Pesquisa em Malária, Instituto Oswaldo Cruz, Fiocruz.* Av. Brasil 4365, Manguinhos, Rio de janeiro. CEP 2104-360, RJ Brazil. These authors contributed equally to the work (Luciana Pereira de Sousa Vieira, Flávia Lima Ribeiro-Gomes). Programa de Pós-Graduação em Ciências Biológicas, Instituto de Ciências Exatas e Biológicas, Universidade Federal de Ouro Preto, Minas Gerais, Brazil (Roberto Farina de Almeida).

## Abstract

Elements of the immune system are necessary for healthy neurocognitive function, and the pattern of the immune response triggered by different exogenous stimuli may induce regulatory or deregulatory signals that can affect nervous functions. Here we investigate the effect of immune stimulation on behavioural parameters in healthy mice and its impact on cognitive sequelae resulting from non-severe experimental malaria. We show that the immune modulation induced by a specific combination of immune stimuli, classically described as capable of inducing a major type 2 immune response, can improve the long-term memory of healthy adult mice and prevent the negative cognitive-behavioural impairments caused by a single episode of mild *Plasmodium berghei* ANKA malaria. This finding has implications for the development of immunogens as cognitive adjuvants.

## INTRODUCTION

The immune and nervous systems may be categorized as plastic cognitive systems due to their ability to recognize real world objects, including microbes, and to their ability to adapt through experience. Following antigenic or sensory stimulation, vertebrate organisms undergo changes in the cellular connections of their immune and nervous systems that alter their abilities and structures. There is considerable evidence for the existence of strong interactions between these two systems^1-7^. Immunomodulation of the nervous system can occur through either physiological or pathological mechanisms. The maturation and homeostasis of nervous cognitive abilities require the participation of components of the immune machinery^6-7^. Exogenous immune stimuli may also have positive or negative effects on the nervous system, depending on the nature and intensity of the immune response elicited^1-4,6^.

Studies on the effects of immune stimuli on brain function have found evidence for i) maternal immune stimulation impairing the neurocognitive performance of offspring^8-9^, ii) both beneficial and harmful effects of neonate vaccination on neuronal plasticity and cognitive function in adulthood^10^, iii) the damaging impact of systemic inflammatory stimuli on the cognitive function of adult mice^11-12^, and iv) neurocognitive dysfunction in both human and experimental models of some infectious diseases^13-27^.

Cerebral malaria (CM), the most severe complication of malaria caused by *Plasmodium falciparum*, can result in neurocognitive sequelae, including motor deficits, behavioural alterations and severe learning difficulties^15^. Long-term negative effects are more common in Africa where the prevalence of *falciparum* malaria and CM is higher^28^. Some of these sequelae are also observed in *Plasmodium berghei* ANKA (*Pb*A) infected C57BL/6 mice, a well-studied model of experimental CM (ECM)^27.^ In recent years, cognitive impairment, mainly related to learning and memory, has also been reported in residents of endemic regions presenting with non-severe malaria^29- 31^. This phenomenon has also been observed in non-severe malaria infections in mice^32^, in which the ECM model was adapted to assess the neurocognitive alterations that occur following a short-term episode of non-severe malaria. Using this adapted model, here we evaluate the effects of immune stimuli on behavioural paradigms such as memory and anxiety, following a mild malaria episode or during homeostasis.

Given the known effect of the immune system on neurocognitive functions, we hypothesized that immune stimulation may affect cognitive performance. Our results show a beneficial effect of immune stimulation on cognitive-behavioural parameters in healthy mice and a reversal of the cognitive impairment caused by malaria parasite infection.

## RESULTS

### Type 2 immune stimuli improve long-term memory in healthy mice

To study the effect of immune stimuli on behavioural paradigms, immunogens were chosen according to the pattern of immune response induced. Three immune stimulation strategies were used: T1 and T2 strategies employed well-known antigens able to induce type 1 and type 2 immune responses, respectively^33-47^, and a “Pool” strategy was created by the combination of T1 and T2 strategies, described in further detail in the Material and Methods section. Briefly, mice were infected with *Plasmodium berghei* ANKA, treated from the fourth day after infection on for seven days, and allowed to rest for thirteen days before being immune stimulated with different strategies (Fig. 1).

**Fig. 1.**
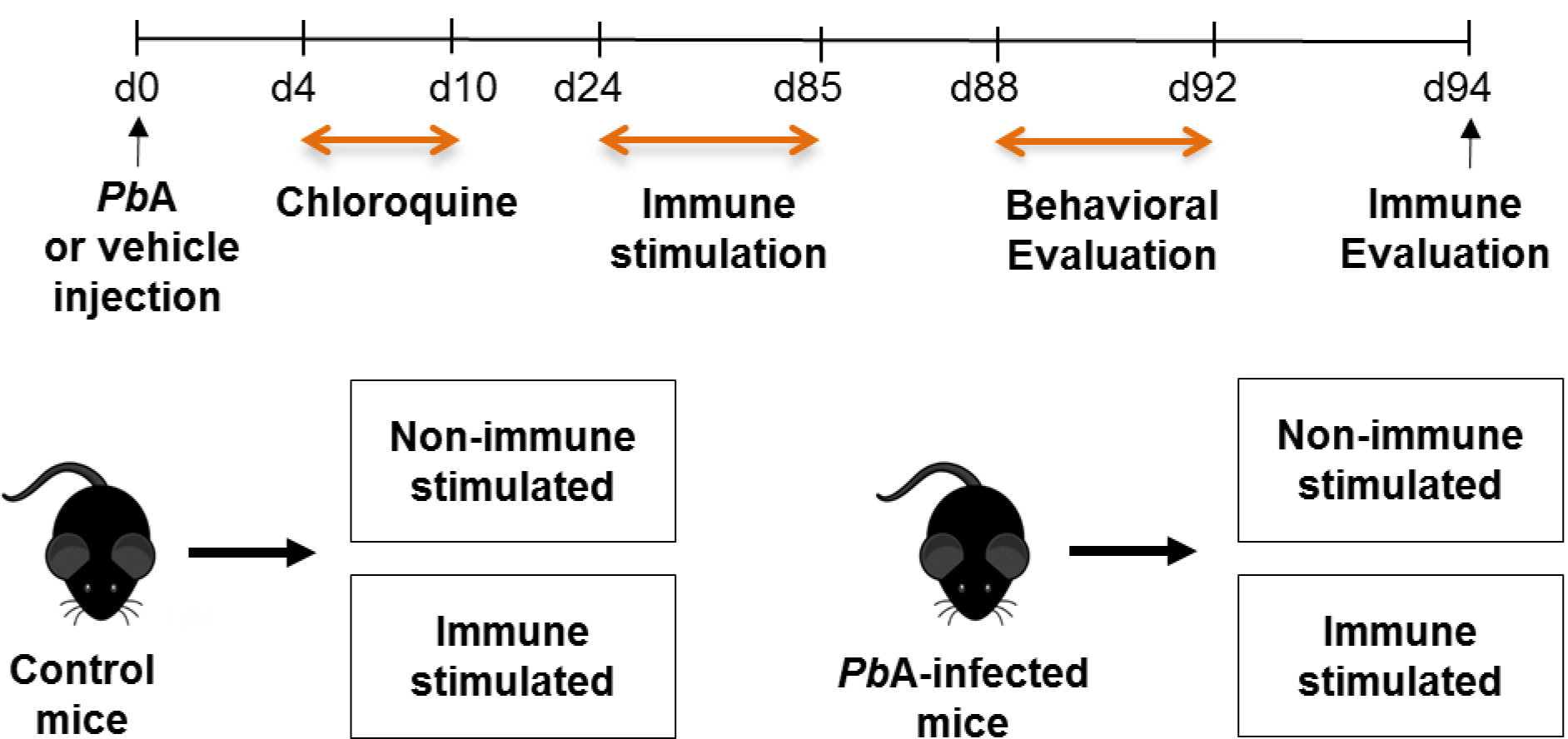
Groups of mice were infected or not with *Plasmodium berghei* ANKA (*Pb*A) and treated with chloroquine (25 mg / kg) for seven days via gavage from the fourth day post-infection. After 14 days, the animals were subdivided into groups of mice immune stimulated with different immunization strategies or non-immune stimulated. Subsequently, mice were evaluated in behavioural tasks for locomotivity, memory and anxiety phenotype. The immune response of mice randomly chosen was evaluated.

The effects of immune responses on locomotion and long-term spatial habituation were assessed via established protocols^32^ in mice subjected to two different sessions of the open field task (OFT), with training (OF1, 10 min.) and test (OF2, 10 min.) sessions 24 hours apart. At the training session, a high rate of locomotor activity is commonly observed. Surprisingly, mice immune stimulated with Pool or T1 strategies showed reduced total OF1 locomotion when compared to non-immune stimulated mice (Extended data, Fig. 1a).

Commonly, after the training session [first OFT (OF1), exposure], exploratory behaviour decreases as the stress related to novelty disappears, and is usually significantly lower after 10 minutes of task performance^32,48-49^. Both non-immune stimulated (Control group) and immune stimulated (Pool, T1 and T2 groups) mice displayed decreased locomotion in the test session (OF2) compared to the training session (OF1) (Extended data, Fig. 1a), as expected. These results indicate that immune stimulation did not affect long-term habituation memory.

Twenty-four hours later mice were subjected to the novel object recognition test (NORT) in the same open field arena. During the training session, a similar exploratory activity of familiar objects (FO1 and FO2) is expected and was observed in all groups of mice (Control, Pool, T1 and T2) (Fig. 2a; Extended data, Fig. 2a), with a mean exploration of 25 seconds (data not shown). Remarkably, mice immune stimulated with the Pool or T2 strategies presented significantly higher recognition memory performance in relation to the Control group during the test session, performed 24 hours later. Mice submitted to the T1 strategy did not differ from the Control group (Fig. 2c; Extended data Fig. 2c). These data indicate that immune stimulation with immunogens that induce type 2 immune responses may enhance long-term recognition memory in healthy mice.

**Fig. 2.**
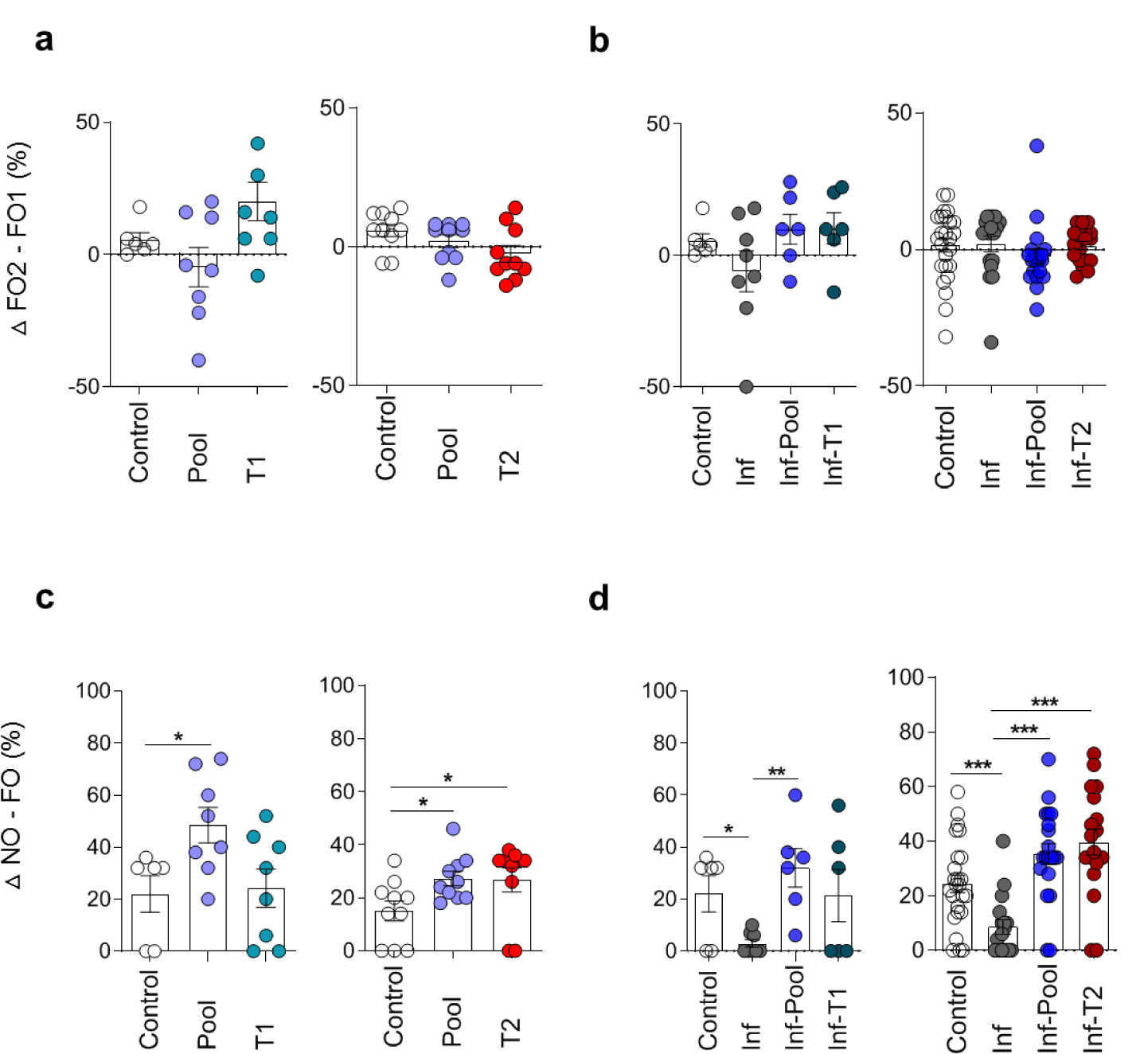
Immune stimulation improves long-term memory performance in healthy and *Pb*A-infected mice. Healthy or *Pb*A-infected (and treated) mice were immune stimulated, or not, with the Pool, T1 or T2 strategies. Behavioural tasks were performed from day 88 to 92 post-infection (77 to 81 days after CQ treatment). During the training session of the New Object Recognition Task (NORT), all experimental groups explored the two objects, called familiar objects (FO1 and FO2, **a, b**) for the same length of time. In the test session, a new object (NO) is introduced in the task (**c, d**). Immune stimulation of healthy mice with Pool and T2 strategies (Pool and T2 groups) improved the exploratory time spent in the NO in relation to the Control group (**c**). *Pb*A-infected mice (Inf group) presented similar exploration of NO and FO, showing a memory deficit, which was reversed after immune stimulation with Pool and T2 strategies (Pool and T2 groups) (d). Experimental groups: Control (non-infected / non-immune stimulated mice, n = 6 - 25); Pool (non-infected / Pool-immune stimulated mice, n = 8 - 10); T1 (non-infected / T1-immune stimulated mice, n = 8); T2 (non-infected / T2-immune stimulated mice, n = 10); Inf (infected / non-immune stimulated mice, n = 6 - 17); Inf-Pool (infected / Pool-immune stimulated mice, n = 6 - 20); Inf-T1 (infected / T1-immune stimulated mice, n = 6); Inf-T2 (infected / T2-immune stimulated mice, n = 8 - 18). Data are expressed as mean and s.e.m. ***P < 0.001; **P < 0.01; *P < 0.05; Mann-Whitney Unpaired t-test was used. Data shown represent one of two to five independent experiments (Control, Pool, T1, T2, Inf, Inf-Pool, Inf-T1); and a pool of two independent experiments (Control, Inf, Inf-Pool, Inf-T2).

### Immune stimulation of healthy mice did not generate an anxiety-like state

In addition to exploratory activity, the OFT also allows the evaluation of phenotypes related to anxiety-like behaviour through analysis of the dwell time or the locomotion rate in the centre of the open field arena during the first exposure to the apparatus. Immune stimulated mice (Pool, T1 and T2 groups) showed no difference in dwell time (data not show) but presented significantly reduced locomotion in the centre of the open field arena in relation to the non-immune stimulated mice (Control group) (Fig. 3a). It seems, however, that this observation may have been influenced by the total reduced locomotion observed in animals submitted to Pool and T1 strategies (Extended data, Fig. 1a). Since no conclusion about anxiety-related behaviour can be confidently extrapolated from these data, we used the light-dark specific task, a conflict avoidance test, to address this issue. In this test, immune stimulated mice (Pool, T1 and T2 groups) clearly behaved similarly to mice of the Control group, remaining an equal time in the light zone (Fig. 3c), and thus implying that immune stimulation did not generate an anxiety-like state.

**Fig. 3.**
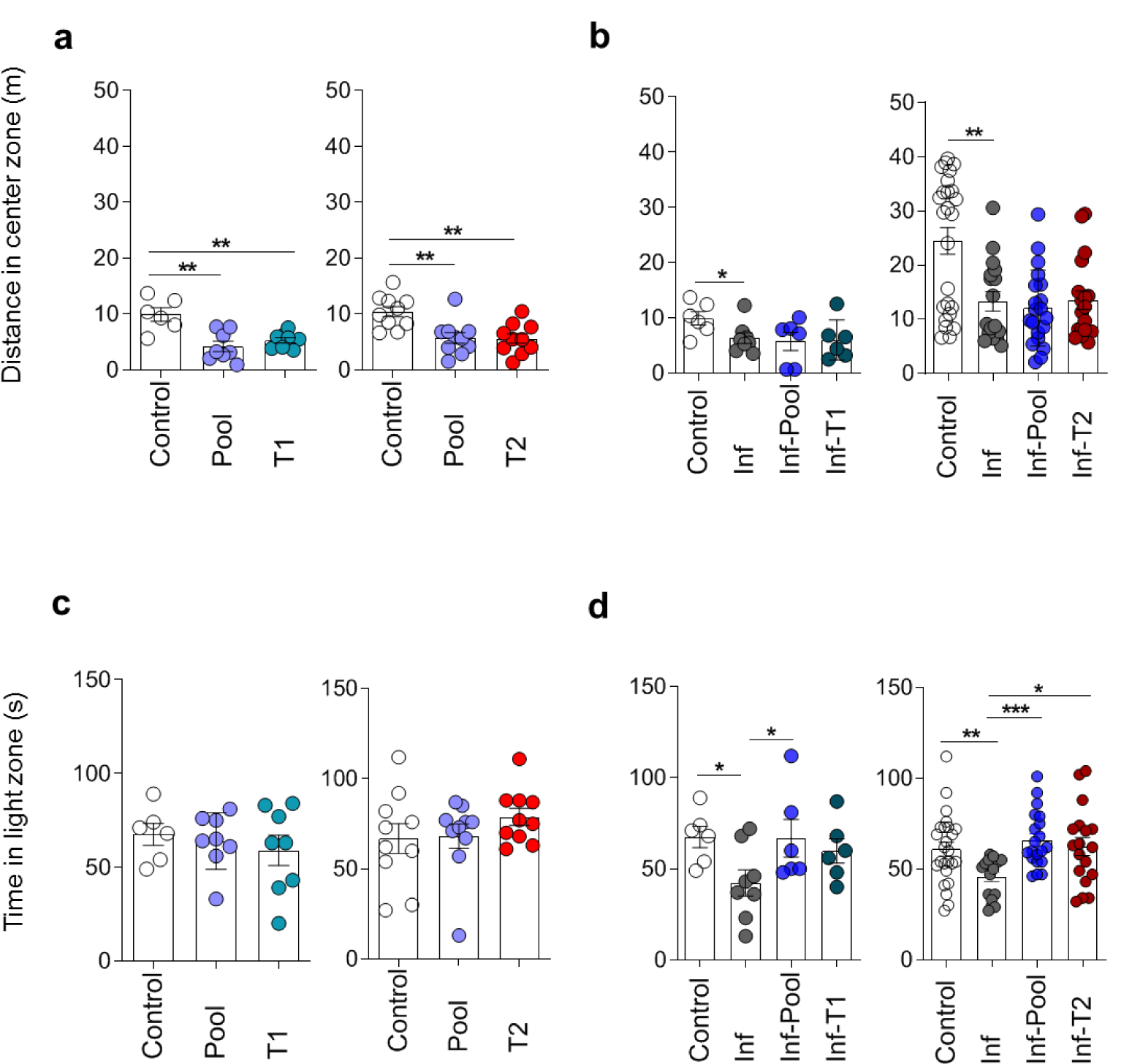
Immune stimulation attenuates the anxiety-like behaviour observed in *Pb*A-infected mice. Healthy or *Pb*A-infected (and treated) mice were immune stimulated, or not, with the Pool, T1 or T2 strategies. Behavioural tasks were performed from day 88 to 92 post-infection (77 to 81 days after CQ treatment). During the training session of the Open Field Task (OFT), Pool, T1 and T2 immune stimulated groups (**a**) and *Pb*A-infected animals (Inf group) (**b**) showed a decrease in the distance travelled in the centre of the arena, as compared to the Control group. No difference was observed between the performance of Pool, T1, T2 and Control groups in the Light / Dark task (**c**). *Pb*A-infected mice (Inf group) spent less time in the light zone of the Light / Dark apparatus. This anxiety-like behaviour was suppressed following immune stimulation with the Pool and T2 immune strategies (**d**). Experimental groups: Control (non-infected / non-immune stimulated mice, n = 6 - 25); Pool (non-infected / Pool-immune stimulated mice, n = 8 - 10); T1 (non-infected / T1-immune stimulated mice, n = 8); T2 (non-infected / T2-immune stimulated mice, n = 10); Inf (infected / non-immune stimulated mice, n = 6 - 17); Inf-Pool (infected / Pool-immune stimulated mice, n = 6 - 20); Inf-T1 (infected / T1-immune stimulated mice, n = 6); Inf-T2 (infected / T2-immune stimulated mice, n = 8 - 18). Data are expressed as mean and s.e.m. ***P < 0.001; **P < 0.01; *P < 0.05; Mann-Whitney Unpaired t-test was used. Data shown represent one of two to five independent experiments (Control, Pool, T1, T2, Inf, Inf-Pool, Inf-T1); and a pool of two independent experiments (Control, Inf, Inf-Pool, Inf-T2).

### Exposure to type 2 immune stimuli may reverse cognitive-behavioural damage caused by non-severe *P. berghei* ANKA infection

About 92% of the world’s malaria cases are due to *Plasmodium falciparum*, 1 to 2% of which progress to cerebral malaria. Therefore, about 90% of all malaria cases globally are caused by this lethal species of *Plasmodium* and occur without apparent clinical complications^28^. Despite the apparent ‘non-severe’ nature of these cases, there is growing evidence that non-severe malaria may impair the cognitive development of children^29-31^.

The experimental model we have previously described uses *Pb*A-infected C57BL/6 mice treated at day 4 post-infection, prior to the appearance of the clinical signs of CM. In our opinion, the main advantage of such a model is that it best mimics the human situation described above that corresponds to the large majority of malaria cases in the world; non-severe falciparum malaria with timely treatment^28^. Using this model, we have been able to observe a long-term cognitive-behavioural impairment related to memory and anxiety as late as 82 days after the end chloroquine (CQ) treatment, when no parasites are present tin the blood^32^.

Given the beneficial effect of immune stimulation on long-term memory in healthy mice described above, we evaluated the effect of the same immune stimuli in mice with behavioural alterations caused by non-severe malaria infection. *Pb*A-infected and treated mice (from here on referred to as the “Infected group”), did not display reduced total locomotion in the training session of the OFT when compared to healthy mice (Extended data, Fig. 1b). However, infected and immune stimulated animals (Inf-Pool and Inf-T2 groups) showed a significant reduction in locomotion in the OF1 when compared to healthy mice (Extended data, Fig. 1b). Control, infected and infected-immune-stimulated groups (Inf-Pool and Inf-T2, but not Inf-T1) displayed normal behaviour with a significant decrease in locomotion in the test session as compared to the training session of the OFT (Extended data, Fig. 1b).

As expected, there was no object preference in the NORT training session since all mice explored both familiar objects for the same length of time (for a mean of 25 seconds; data not show) (Fig. 2b, Extended data, Fig. 2b). Consistently, infected mice presented long-term recognition memory sequelae that manifested as similar exploration of the familiar object (FO) and new object (NO) in the NORT. This impairment disappeared following stimuli with Pool or T2 immunization (Fig. 2d, Extended data, Fig. 2d), pointing to a beneficial effect of immune stimulation triggered by type 2 immunogens in reversing of the cognitive deficits associated with malaria.

### *P. berghei* ANKA infection in mice induces an anxiety-like behaviour that is reversed by immune stimulation with type 2 immunogens

The distance travelled in the periphery and in the centre of the open field arena are inversely related. Since the latter was decreased in *Pb*A-infected mice (Fig. 3b) and no change in the locomotion during the training session (OF1) occurred among Control and Infected groups (Extended data, Fig. 1b), the decrease may be interpreted as the expression of an anxiety-like behaviour. This behaviour was confirmed by the observation of a reduction in time spent, by infected mice, in the light zone of the light-dark task, a more sensitive and widely used test to evaluate anxiety-related parameters in rodent. The anxiety-like behaviour was reversed by Pool and T2, but not by T1, strategies of immune stimulation (Fig. 3d).

### Immune stimulation procedures and non-severe *P. berghei* ANKA malaria elicit immune responses

The specific immune responses triggered by the immunogens in the Pool, T1 and T2 strategies (tetanus toxoid, influenza, *Pf*MSP3 and OVA) were evaluated at the end of the behavioural task experiments, and the effectiveness of the stimuli was confirmed (Extended data, Fig. 3a,b,c,d). No specific humoral immune response was observed against diphtheria toxoid (data not show), confirming previous observations of the low immunogenicity of diphtheria toxoid in mice compared to other experimental models^50^.

At the time the immune responses were evaluated (84 days after the end of CQ treatment), non-immune stimulated infected animals did not present increased levels of serum cytokines when compared to the Control group (Extended data, Fig. 4a,b,c,d). However, higher levels of TNFα, IFNγ, IL-6, IL-10 and/or IL-4 were detectable in all groups of mice stimulated with T1, T2, or Pool strategies (Extended data, Fig. 4a,b,c,d), ratifying the immune stimulation by the different strategies used. Interestingly, only IL-10 was consistently increased among healthy and infected mice stimulated with Pool or T2 strategies, although statistical significance was not achieved between Pool and Control groups (Extended data, Fig.4e).

We evaluated the splenic immune response of healthy and infected mice exposed to Pool and T2 strategies, since only these approaches were able to immunomodulate the cognitive behaviour of mice. The immune stimulated healthy mice showed increased spleen weight and total number of splenocytes (Extended data, Fig. 5a,b,c). The weight and total number of splenocytes in Infected animals were not different to those in the Control group (Extended data, Fig. 5a,b,c). As observed in healthy immune stimulated animals, immune stimulation of *Pb*A-infected mice *via* Pool or T2 strategies induced splenomegaly (Extended data, Fig. 5a,b,c).

Healthy mice immune stimulated with either Pool or T2 strategies presented similar patterns of modulation of different immune components. We observed an increase in the frequency of splenic B cells (Extended data, Fig. 6b), CD4 and CD8 T cells with central memory phenotype (Extended data, Fig. 7d,g) and CD4 T cells with regulatory function (Treg cells) (Extended data, Fig. 6e) in both Pool and T2 immune stimulated groups when compared to non-immune stimulated animals. A reduction in the frequency of CD8 T cells was also observed in mice immune stimulated with the T2 strategy when compared to the Control group (Extended data, Fig. 6d).

*Pb*A-infected mice had higher frequencies of B cells, total CD4 and CD8 T cells (Extended data, Fig. 6a,b,c), and CD4 and CD8 T cells with naïve and central memory phenotypes (Extended data, Fig. 7a,b,e,d,g) when compared to healthy mice (Control group). The frequency of Treg cells, however, was similar between infected and healthy mice (Extended data, Fig. 6e).

Immune stimulation of *Pb*A-infected mice with Pool or T2 strategies induced comparable increases in the frequencies of splenic B cells, Tregs (Extended data, Fig.6a,b,e), effector/effector memory CD4 T cells and central memory CD8 T cells (Extended data, Fig.7a,c,g), and reduction in the frequencies of total CD8 T cells when compared to non-immune stimulated infected mice (Extended data, Fig. 6a,d).

In summary, immunological analysis demonstrated that, independently of the health status of the mice, immune stimulation with type 2 immunogens reduces the frequency of CD8 T cells and increases the percentage of Treg cells in the spleen, as well as the serum level of IL-10.

Taken together, our data point to a positive influence of immune responses induced by strategies involving type 2 stimuli on the long-term memory of healthy mice, confirm our previous demonstration of late neurocognitive behavioural dysfunction following a single episode of non-severe malaria, and indicate a recovering effect of this deficit exerted by immune stimulation with type 2 immunogens subsequent to infection.

## DISCUSSION

Here, we describe for the first time a beneficial modulatory effect of immune stimulation on cognition in healthy adult mice. Our findings show a clear positive effect of immune stimuli, specifically triggered by immunization strategies involving type-2 immunogens, on long-term memory, as verified by the ‘new object recognition task’ (NORT), a robust and frequently used behavioural task for the analysis of recognition memory in mice^51^.

We have previously identified cognitive-behavioural impairment as late sequelae of a single non-severe malaria episode, using the classical ECM model with treatment of animals before the presentation of neurological signs or cerebrovascular damage^32,52^. We propose that this model is appropriate for the study of non-severe *Plasmodium falciparum* malaria^28^, as both parasite-host pairs involve the potentiality of CM development that can be avoided with timely drug treatment.

The data described here confirm our previous work, showing that neurological impairment can occur even in the absence of classical clinical signs of CM^32^. We propose, therefore, that the term “non-severe malaria” should be used, preferentially to the classical expression “non-cerebral malaria”, to describe the experimental model or the human situation in which clinical signs of CM are not observable. In agreement with our observation is the activation of microglia at day 4 post-infection, before the overwhelming cerebral inflammation and development of the clinical signs of CM^53^. The levels of proinflammatory cytokines also increase around 3-4 days after *P. berghei* ANKA infection in C75BL/6 mice^54,55^. It is possible, therefore, that the late cognitive deficit observed in our studies results from the early activation of immune cells in the central nervous system (CNS).

Remarkably, we observed a positive effect of immune stimulation on reversing the cognitive-behavioural impairment associated with non-severe malaria. Mice treated with CQ four days after infection by *P. berghei* ANKA and immune stimulated with T2 and Pool strategies did not present the deficit of object recognition recorded after infection without subsequent immune stimulation. We also observed reversal of anxiety-like behaviour in a light-dark task, following immune stimulation of infected mice. Recent data from our laboratory shows that these behavioural changes are observable as early as 12 days subsequent to malaria treatment (data not shown), pointing to a reversible potential effect of the immune stimuli.

The CNS and the immune system interact under homeostatic conditions and a well-balanced immune response is needed for a proper function of the CNS^6-7^. T cells are essential for normal neurogenesis and cognition^6,56-58^.

Communication between peripheral immune cells and CNS takes place in the brain, probably at the meningeal spaces^6^, where T cells influence the CNS via the production of cytokines. It has been shown that proinflammatory cytokines impair cerebral function and cognition at high pathological concentration, as during infections^6^. An exacerbated peripheral inflammatory response may cause M1 microglial activation and provoke the production of proinflammatory cytokines such as TNF-α and IL1-β that may impair cognitive function^59^. Elevated levels of anti-inflammatory/regulatory cytokines such as IL-4 and IL-10 may have the opposite effect, inducing M2 microglial activation and positively influencing cognition^60-62^.

Treg cells are a subset of T cells with immunomodulatory function, important for immune and neuronal homeostasis under physiological conditions, and for the control of pathological immune responses^63-66^. They perform their function mainly *via* secretion of IL-10 and TGFβ, anti-inflammatory/regulatory cytokines^63-66^. After ischemic brain stroke, there is massive accumulation of Treg cells in the mouse brain^67^, where they decrease inflammatory cell infiltration and microglia activation, antagonize the production of proinflammatory cytokines and, consequently, reduce brain damage through a mechanism involving IL-10 secretion^68^. The neuroprotective activity of Treg cells has also been described in murine models of Parkinson’s disease, HIV-1-associated neurodegeneration and amyotrophic lateral sclerosis^69-72^.

In this study, healthy and infected-mice stimulated with strategies involving type 2 immunogens (Pool and T2 groups) significantly increased the number of splenic Treg cells and IL-10 level in the serum. Considering that Treg cells and IL-10 can restrict neuroinflammation^71-76^, it is reasonable to assume that the immunization strategies used likely improve cognitive function by promoting a balanced cross-talk between the immune system and the CNS mediated through Treg cells and IL-10. The mechanism by which immune stimulation with type-2 immunogens benefits cognition is presently under investigation.

The results reported here may offer a new paradigm for the design of memory improvement strategies. Our data suggest that vaccination procedures may provide benefits additional to the prevention of infection, offering a potential approach for boosting cognition function in healthy individuals, and in helping the recovery of those whose cognition may have been impaired by chronic and infectious diseases, including malaria, and by the effects of ageing.

## ACKNOWLEDGEMENTS

LPS is grateful to the *Programa de Pós-Graduação em Biologia Parasitária* of *Instituto Oswaldo Cruz (IOC), Fundação Oswaldo Cruz (Fiocruz)* for the Doctoral fellowship. The authors are grateful to Professor Richard Culleton for his kindness carefully reading the final version of this manuscript and for the valuable comments and suggestions for the improvement of the text. We do also thank Doctor Leonardo Carvalho for his critical review and welcome discussions. We are indebt to the *Laboratório de Inflamação* (Dr. Marco Aurélio Martins and Dr. Tatiana Ferreira), *Laboratório de Pesquisa em Malária* (Luana santos and Thalita Ferraz), *Laboratório de Pesquisa sobre o Timo* of the *IOC-Fiocruz* (Dr. Daniella Mendes Arêas and Dr. Dyna Raposo) of *IOC; Biomanguinhos* (Dr. Maria de Lourdes de Sousa Maia, Alessandro Fonseca, Camilla Bayma and Dr. Denise Cristina Matos) and *Farmanguinhos* (Dr. Márcia Coronha Ramos Lima and Dr. Andréa Luca) of *Fiocruz and Instituto Butantan* (Dr. Jorge Kalil and Dr. Paulo Lee Ho and Aline Abrantes) and Vac4all (Dr. Pierre Duiilhe) for reagent supply and study of the immune responses to vaccines.

## AUTHOR CONTRIBUTIONS

LPS was responsible for the realization of all experiments (including infection and treatment; immune stimulation, and conduction, observation and data collection/systematisation of cognitive tests and immune response analyses in mice), helped in the analysis and interpretation of tests and drafted the manuscript. FLRG followed all stages of the experiment, realization of experiments, discussed the protocols and the project, was in charge of the analysis and discussion of immune response data and helped in drafting the manuscript. RFA and TMS helped in systematization of data concerning behavioural tests and analysed and interpreted the cognitive data. GW proposed the statistical analyses of the data and was responsible for them. DOS discussed the project since its conception and helped in designing the experiments. CTDR is responsible for conception and design of the study, and helped in data analysis, interpretation and drafting and finalizing the manuscript together with LPSV and FLRG. All authors read, reviewed and approved the final manuscript.

## METHODS

### Mice and Parasite

The *Instituto de Ciência e Tecnologia em Biomodelos* of the *Fundação Oswaldo Cruz* (ICTB-*Fiocruz*, Brazil) provided seven-week-old C57BL/6 female mice weighing 20-25 g. Mice were housed in racks with an air filtration system in a room maintained at 25°C and light/dark cycles of 12 hours in cages containing five animals with free acquisition to food and water. All procedures were carried out in accordance with animal welfare approved by the Ethical Committee on the Use of Laboratory Animals of *Instituto Oswaldo Cruz* under *CEUA-IOC*: L-010/2015 concession. *Plasmodium berghei* ANKA (*Pb*A) infections were carried out using a stable transfected strain of *Pb*A expressing a green fluorescent protein (*Pb*A-GFP) generated as described previously^1^.

### Infection and treatment of experimental groups

C57BL/6 mice were infected intraperitoneally (ip) with 150 μl of *Pb*A-infected red blood cells, cryopreserved and thawed. Five days after infection, the total blood was collected, adjusted to 1 ⨯10^6^ parasitized erythrocytes in 100 μl of PBS and injected ip to C57BL/6 mice from the experimental groups. Parasitaemia was monitored by flow cytometry, based on the percentage of GFP^+^ erythrocytes. In this experimental model, the establishment of cerebral malaria (CM) occurs between the fifth and sixth day of infection^2^. In this study, mice were treated on the fourth day of infection (mean parasitaemia 2.5%) with 25 mg/kg of chloroquine (CQ) by gavage for seven days^3^, before any clinical sign of CM. All groups were similarly manipulated. Experiments carried out with groups of uninfected mice treated with CQ or not (control group received PBS) have previously shown that the CQ treatment did not influence the performance in behavioural tasks and anxiety phenotype^4^.

### Experimental Description

C57BL/6 mice were divided into groups of *Pb*A-infected and Control animals (non-infected) and both were treated with chloroquine (CQ) for seven days from the fourth day of infection. Thirteen days after treatment, mice from respective groups were subdivided into non-immune stimulated and immune stimulated groups (Fig. 1). The following vaccines and antigens were used for immune stimulation: Diphtheria and Tetanus toxoids (dT) vaccine for adults, Influenza vaccine, *Plasmodium falciparum* Merozoite Surface Protein 3 (*Pf*MSP-3 recombinant protein), White chicken egg ovalbumin (OVA) and Lipopolysaccharide of *Escherichia coli* (*Ec*LPS). Three different immune stimulation strategies were performed: a combination of all antigens and vaccines described above (from here now, called Pool); a combination of antigens and vaccines (Influenza vaccine and *Ec*LPS) that trigger, preferentially, a type 1 pattern of immune response (from here now, denominated T1); and a combination of antigens and vaccines (dT vaccine, *Pf*MSP-3 recombinant protein and OVA) that trigger, preferentially, a type 2 pattern of immune response (from here now, called T2). The groups of mice were denominated: Control (non-infected / non-immune stimulated); Pool (non-infected / Pool-immune stimulated); T1 (non-infected / T1-Immune stimulated); T2 (non-infected / immune stimulated); Inf (infected / non-immune stimulated); Inf-Pool (infected / Pool-immune stimulated); T1 (infected / T1-immune stimulated) and T2 (infected / T2-immune stimulated). All control groups were treated as the experimental groups: they were age-matched, mock-immune stimulated, mock-infected, and treated with CQ whenever appropriate. Subsequently, mice behavioural performance was assessed by Open Field Test (OFT), Novel Object Recognition Test (NORT) and Light-Dark (Fig. 1). About 300 mice were used for these experimental strategies in five consecutive sessions.

**Fig. 1.**
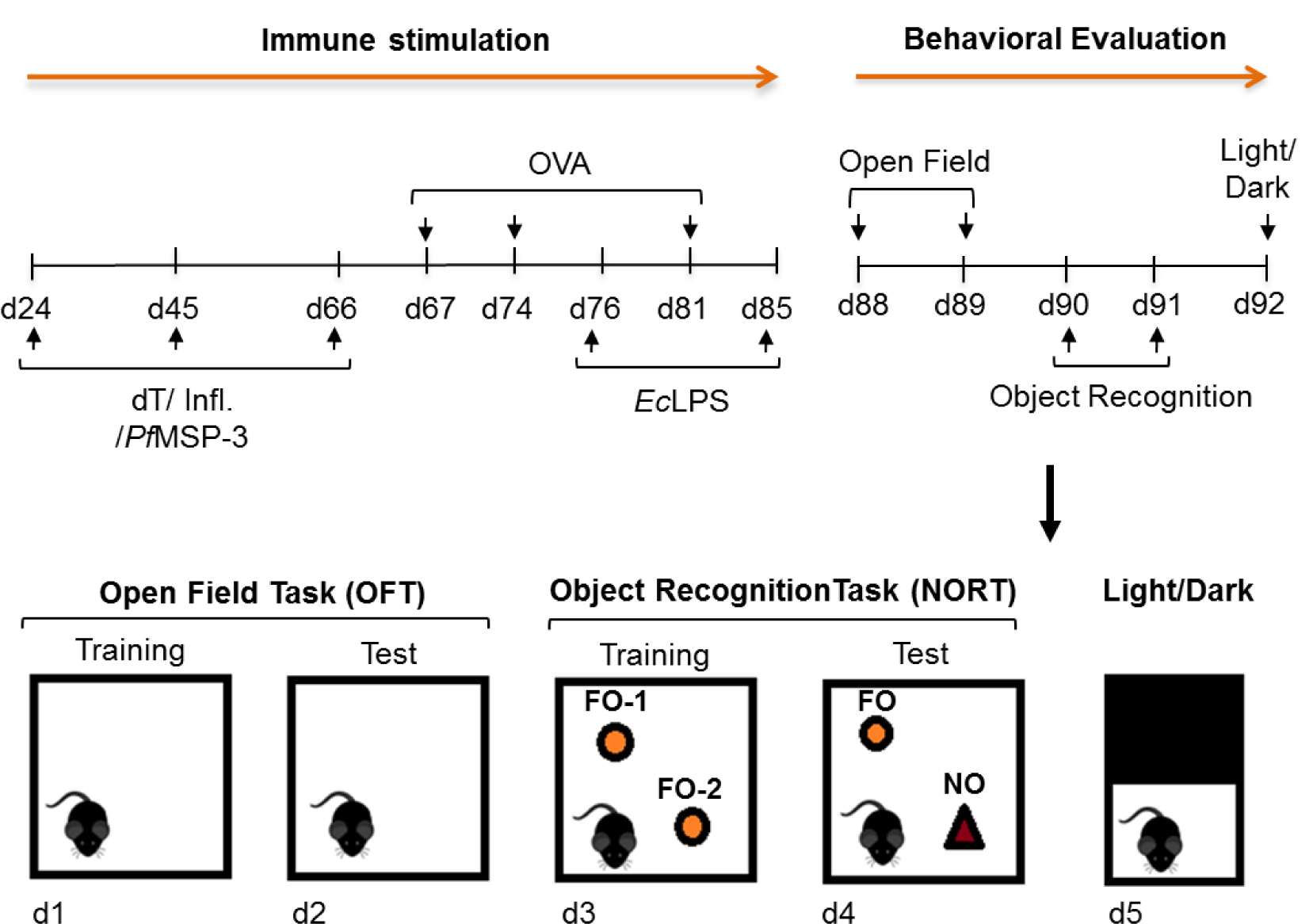
Material & Methods. Flow chart of immune stimuli and behavioural assessment. Mice (infected or not with *Pb*A and treated with CQ) were immune stimulated or non-immune stimulated, according to the composition of the immunization (Pool, T1 and T2) strategies used. Three doses of the dT and Influenza vaccines and the *Pf*MSP-3 recombinant protein were inoculated conjointly, in different pathways, with a twenty-day interval between inoculations. Three doses of OVA, with a six-day interval between each one, were inoculated one day after the third dose of dT and Influenza vaccines and *Pf*MSP-3 protein. The first injection of *Ec*LPS was done two days after the second dose of OVA, being the second of two injections administered nine days after the first one. Assessment of performance on behavioural tasks started 88 to 92 days post infection (77 to 81 days after the complete parasitological cure of animals obtained with CQ treatment). The beginning of behavioural tests corresponded to 22 days after the last stimulation with the vaccines (Tetanus-Diphtheria and Influenza) and the *Pf*MSP-3 recombinant protein; 7 days after the latter injection of Ovalbumin; and 3 days after the LPS final inoculation. The open field was performed to measure locomotivity, spatial habituation memory and anxiety phenotype, in two sessions [training (OF1) and test (OF2)]. Thereafter, the new object recognition task (NORT) was performed to measure long-term recognition memory, also in two sessions at consecutive days (training and testing). Finally, the anxious behaviour phenotype was specifically evaluated, by the light-dark test, in a unique session.

### Immune system stimuli

The immune stimulation was initiated fourteen days after the end of CQ treatment, being performed in the course of the following sixty-two days (Fig. 1). Antigens and/or vaccines were administered by different routes and in different regions of the animal’s body (Table 1). The doses administrated were defined based on dose-response protocols available in the literature capable of stimulating the murine immune system without imparting risk of death to mice immune stimulated^5-12^.

**Table 1.**
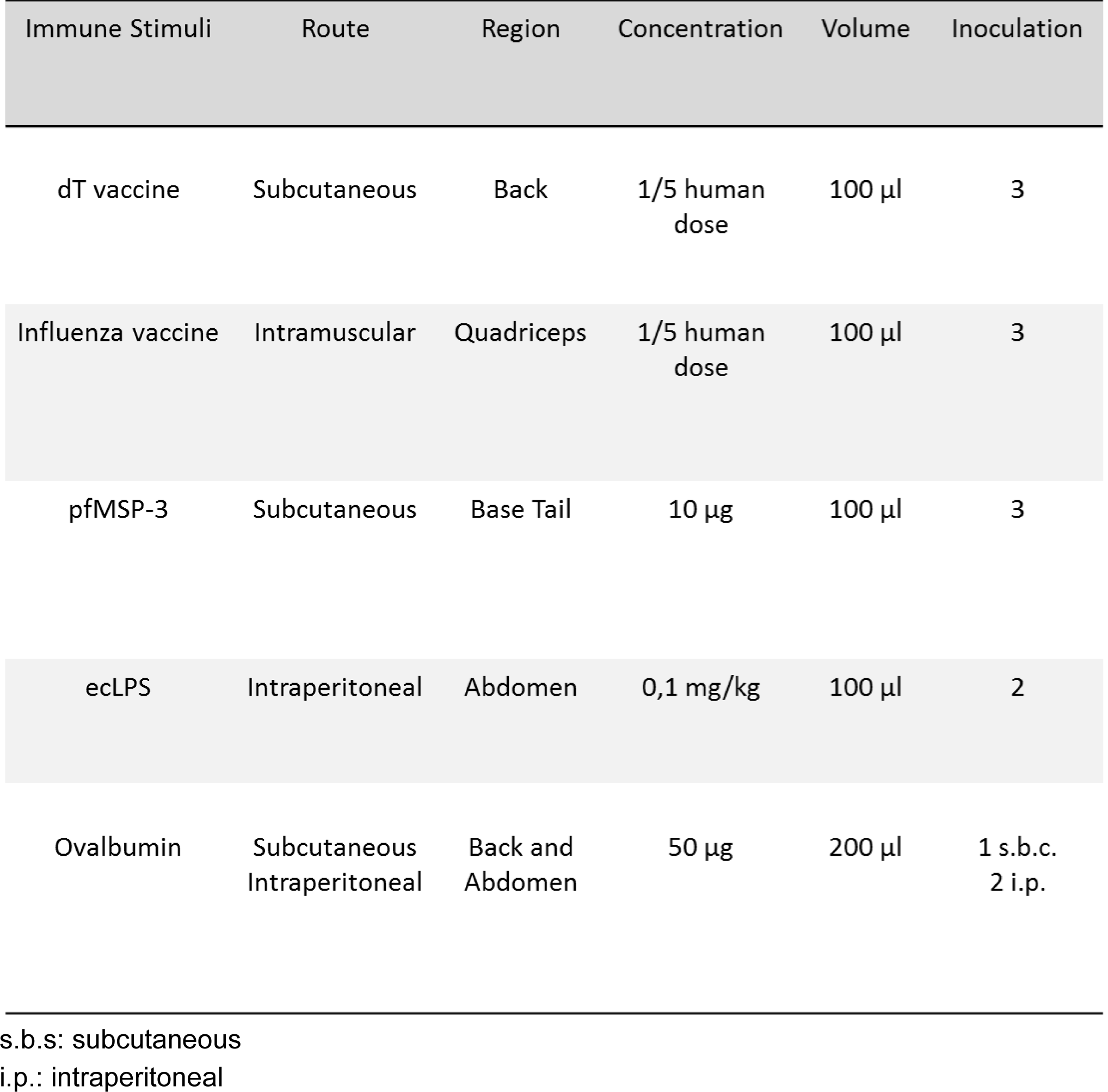
Immune stimulus inoculation strategy: route, region, concentration, volume and number of injections of immunogens.

#### Immune stimulation with Plasmodium falciparum Merozoite Surface Protein 3 (PfMSP-3 recombinant protein)

The mice were challenged with 10 μg of *Pf*MSP-3/mice recombinant protein (in collaboration with Clinical Trials of Malaria Vaccines (Vac4All Initiative), Paris, France) adsorbed on 70% adjuvant solution MONTANIDE™ ISA 50 V2 W / O (SEPPIC. Air Liquide - Healthcare), in 100 μl of PBS. Three subcutaneous injection were performed at the tail region with a twenty-day interval between immune stimulations^9-10^ (Fig. 1, Table 1).

#### Immune stimulation with Tetanus-Diphtheria and Influenza Vaccines

The vaccines used in this study were: Tetanus-Diphtheria (dT) double bacterial (Biological E Limited - BE, Telangana - India, Lot. 34005815), in collaboration with the Division of Health Surveillance - CAP 3.1 of the *Fundação Oswaldo Cruz* (Rio de Janeiro, Brazil); and Trivalent Influenza granted by the Technological Development and Production Division of *Instituto Butantan* (São Paulo, Brazil, Lot. 160034). Mice received 100 μl (1/5 of the human dose) of dT and Influenza vaccines by subcutaneous (dorsal region) and intramuscular (left quadriceps region) routes, respectively (Table 1). Three inoculations with a twenty-day interval between immune stimulations were performed^5-6^ (Fig. 1).

#### Immune stimulation with Ovalbumin (allergen)

Mice received 50 μg/mice of white chicken egg ovalbumin (SIGMA-ALDERICH, Cod. A5503-50g) adsorbed onto aluminum hydroxide [Al (OH) 3] in a final volume of 200 μl per animal in three inoculations. The first inoculation was performed at the dorsal region by subcutaneous injection and the following (second and third inoculation) by ip route (Table 1) with six days between them^11^ (Fig. 1).

#### Immune stimulation with Lipopolysaccharide from *Escherichia coli* (Ec*LPS*)

Mice were challenged with 0.1 mg/kg of *Ec*LPS O111: B4 (SIGMA-ALDERICH, L2630-10MG, Lot 025M4040V12140701) diluted in phosphate-buffered saline (PBS). Two ip inoculations were performed (Table 1) with a range of nine days between the immune stimulations^12^ (Fig. 1).

### Evaluation of the immune response

Following stimulation of the immune system, mice were randomly selected and sacrificed for individual withdrawal of whole blood, *via* cardiac puncture, and spleen at day 84 after the end of CQ treatment. Serum samples were preserved at −70 °C. Total IgG antibody response to *Pf*MSP-3 recombinant protein, Tetanus-Diphtheria toxoids (dT) and Influenza vaccines; the serum cytokine profile; the splenic lymphocyte subpopulations; and the response to Ovalbumin sensitization were evaluated.

#### Specific antibody responses

The antibody response against *Pf*MSP-3 recombinant protein and Influenza vaccine were determinate by conventional Enzyme-Linked immunosorbent Assay – ELISA^9-10^, and the antibody response against dT vaccine was determined by Toxin Binding Inhibition – ToBI^13^.

#### Cytokine profile

Cytokines in the serum samples were measured with Cytometric Bead Array (CBA) Mouse Th1/Th2/Th17 (BD Biosciences) according to the manufacturer instructions. The data were collected on the BD FACSCANTO II flow cytometer and analysed by FCAP Array^**TM**^ Software (BD Bioscience).

#### Splenic lymphocyte subpopulations

Individual spleens were removed and mechanically dissociated using a syringe plunger above 70 μm-pore size Falcon cell strainer (BD Biosciences). Red blood cells were lysed using ACK lysing buffer (Sigma). Single-cell suspensions were counted and incubated with anti-Fcγ III/II (CD16/32) receptor Ab (2.4G2) in PBS containing 3% FCS for 15 min, and immunolabelled for 30 min at 4°C in the dark with the following fluorochrome-conjugated antibodies: PE-Cy7 anti-mouse CD8 (53-6.7), PerCP-Cy5.5 anti-CD3 (145-2c11), APC-H7 anti-mouse CD4 (GK1.5), APC anti-mouse B220 (RA3-6B2), BB515 anti-mouse CD62L (MEL14), APC anti-mouse CD44 (IM7) and/or PE anti-mouse CD25 (7D4). For Treg cells analyses, cells were fixed and permeabilized, after staining for surface markers, with eBioscience™ Foxp3/Transcription Factor Staining Buffer Set according to the manufacturer instructions and incubated with the antibody Alexa Flour 647 anti-Foxp3 (R16715). All antibodies were from BD Biosciences. Data were collected using FACSDiva software on a FACSCANTO II flow cytometer (BD Biosciences), and analysed using FlowJo software (TreeStar).

#### Intradermal skin test

In the footpad of the left paw, 3 μg of OVA, diluted in 30 μl of PBS, were injected in each animal. After 30 minutes, the plantar thickness (mm) was measured using a digital caliper. Oedema formation was expressed as the difference of the pad thickness measured before and after the inoculation of OVA^11^.

### Behavioral analysis

The schedule of the behavioral tasks is shown in Methods Fig. 1. Mice were individually submitted to different behavioral paradigms to evaluate their exploratory and locomotor activity, cognitive abilities, and parameters involved in anxiety-like behavior from day 88 to 92 post-infection (77 to 81 days after the complete parasitological cure of animals obtained with CQ treatment). The beginning of behavioral tests corresponded to 22 days after the last stimulation with *Pf*MSP-3 recombinant protein, Tetanus-Diphtheria, and Influenza vaccines, 7 days after the last injection of ovalbumin, and three days after the LPS final inoculation. The same cohort of mice was used in all tasks (Methods Fig. 1). All experiments were carried out with an incandescent light source of 200 lux of intensity in the evening period. Animals were acclimatized in the experimental room at least for 2 hours before the experimental sessions. Behavior was captured by a video camera positioned above the task apparatus. Locomotion in the open field and the object recognition task was analyzed by the AnyMaze® software (Stoelting Co., Wood Dale, IL, USA), while a trained blind-to-treatment researcher evaluated other behavioral parameters by video analysis. In all behavioral tests, mice were individually placed on the apparatus, which was previously cleaned with 70% alcohol and dried.

#### Open Field Task (OFT)

To address the effect of immune stimuli on locomotion and on long-term habituation, mice were individually submitted to the OFT with a training (OFT1) and a test (OFT2) session 24 hours apart, as described elsewhere^4^. In each OFT session, mice were individually allowed to freely explore a grey acrylic square box, dimensions (50 × 50 × 50 cm, length × width × height), for 10 minutes. In OFT1, locomotor activity was evaluated during the first three minutes (short-term habituation to novelty) and the last six minutes of the session, and the time and distance traveled in the center zone during the entire session. In OFT2, we evaluated the first three and the last seven minutes of the total distance traveled.

#### Novel Object Recognition Task (NORT)

To evaluate long-term memory for object recognition, the NORT was carried out in the OFT apparatus, 24 hours after its OFT2 session^4^. In the training session, mice were exposed to two identical objects, called familiar objects (FO1 and FO2), for which similar exploratory activity was expected^15^, since they were both novel. The test session was carried out 24 hours later when mice were exposed to a new object (NO) and to one of the previously exposed familiar objects (FO1 or FO2). Memory expression is indicated by the tendency of the animal to spend more time exploring the NO rather that the FO^4,15^. Animals were individually placed in the periphery of the box with the objects in a session for 10 minutes. Exploration was recorded only when the animals touched the objects, located in opposite and symmetrical corners of the box, with their nose or mouth. The time of exploration of each object was recorded, and its percentage of the time of exploration of both objects was calculated. The object recognition index is calculated as the percentage of time spent on each object (referred to the total time spent on both objects). The difference between the time spent with the NO and the FO is expressed as a delta value obtained with the subtraction of the indexes of each object.

#### Light/Dark Task

The light/dark task was carried out as described by Almeida *et al*.^16^ with minor modifications to evaluate the anxiety behavior-like phenotype^17^. The apparatus was a rectangular acrylic box (50 x 30 x 30 cm, height × length × width) with two sides colored white and black, separated by a wall (5×5cm) with an opening at the level of the base of the apparatus joining both sides. A white 100W lamp, placed 60cm above the center of the apparatus, illuminated the white side of the apparatus, while the black side was kept closed without illumination. The mice were individually placed in the light compartment for free exploration of the apparatus for 5 minutes. The following behavioral parameters were analyzed: the time spent in the light compartment and the number of transitions between the compartments (light and dark).

### Statistical analysis

All statistical analyses were performed using a statistical software package (Prism 5.0, GraphPad). The data were extracted from the AnyMaze® software. To analyse OFT and Light / Dark task, we used the absolute data. The time in each object in NORT was transformed into a percentage, from which the delta was extracted based on the subtraction: OF1 - OF2 (training session) and NO - FO (test session). The two-way ANOVA with Bonferroni correct were used to analyse OFT. The Student t-test with Mann-Whitney correction were used for to analyse the groups in NORT, light-dark task and immune response. Data are presented as mean ± standard error. P <0.05 was considered statistically significant.

---

**Extended data, Fig. 1.**
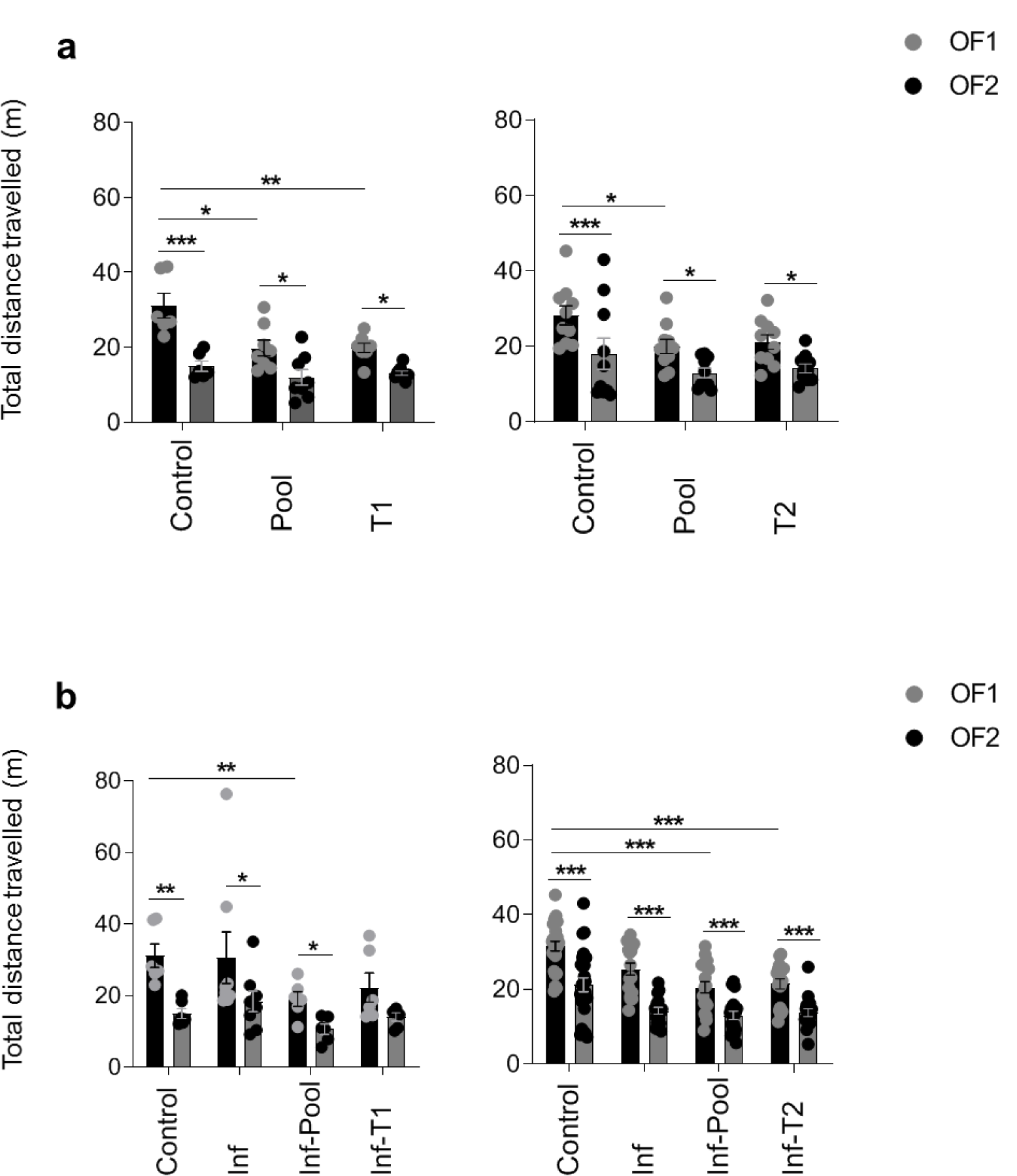
Immune stimulation and *Pb*A infection do not influence habituation memory in mice. Healthy or *Pb*A-infected (and treated) mice were immune stimulated, or not, with the Pool, T1 or T2 strategies. Behavioural tasks were performed from day 88 to 92 post-infection (77 to 81 days after CQ treatment). Total distance travelled in the Open Field Task (OFT) during the training (OF1) and test session (OF2) in healthy and infected mice (**a, b**) were evaluated. OFT: healthy mice groups (Control, n = 6; Pool, n = 8; T1, n = 8 and Control, n = 10; Pool, n = 10; T2, n =10) and infected mice groups (Control, n = 6; Inf, n = 8; Inf-Pool, n = 6; Inf-T1, n = 6 and Control, n = 25; Inf, n = 17; Inf-Pool, n = 20; Inf-T2, n = 18). Data shown represent one of two to five independent experiments (Control, Pool, T1, T2, Inf, Inf-Pool, Inf-T1); and a pool of two independent experiments (Control, Inf, Inf-Pool, Inf-T2).

**Extended data, Fig. 2.**
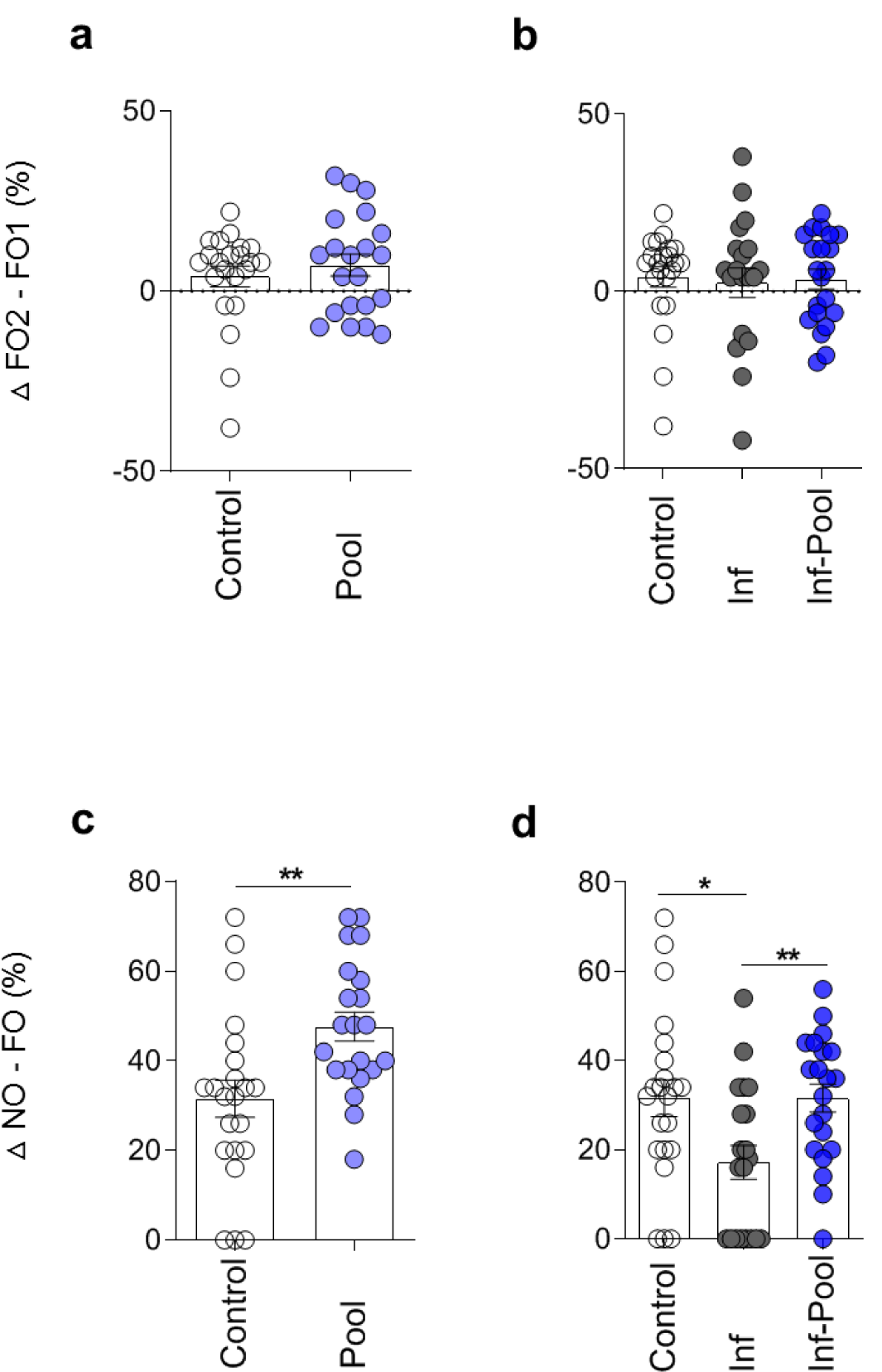
Immune stimulation improves long-term memory performance in healthy and *Pb*A-infected mice. Healthy or *Pb*A-infected (and treated) mice were immune stimulated, or not, with the Pool strategy. Behavioural tasks were performed from day 88 to 92 post-infection (77 to 81 days after CQ treatment). The exploration of the two familiar objects (FO1 and FO2), during the training session of NORT (**a, b**), and of the FO and the novel object (NO), during the test session (**c, d**), were explored and are expressed as differences in percentage of the exploration time. All groups of mice explored similarly the FO1 and FO2 during the training session (**a, b**). Immune stimulation of healthy mice with Pool strategy (Pool group) improved the exploratory time spent on the NO in relation to the FO, as compared to the Control group (**c**). Pool-immune stimulation of *Pb*A-infected mice (Inf-Pool group) reversed the memory deficit of *Pb*A-infected mice (Inf group) (**d**). NORT: healthy mice group (Control, n = 22; Pool, n = 21) and infected mice group (Control, n = 22; Inf, n = 20; Inf-Pool, n = 21). Data shown represent a pool of two independent experiments. Data are expressed as mean and s.e.m. \*\*\**P* < 0.001; \*\**P* < 0.01; \**P* < 0.05; Two-way ANOVA (a, b) and Unpaired t-test (c, d, e, f) was used.

**Extended data, Fig. 3.**
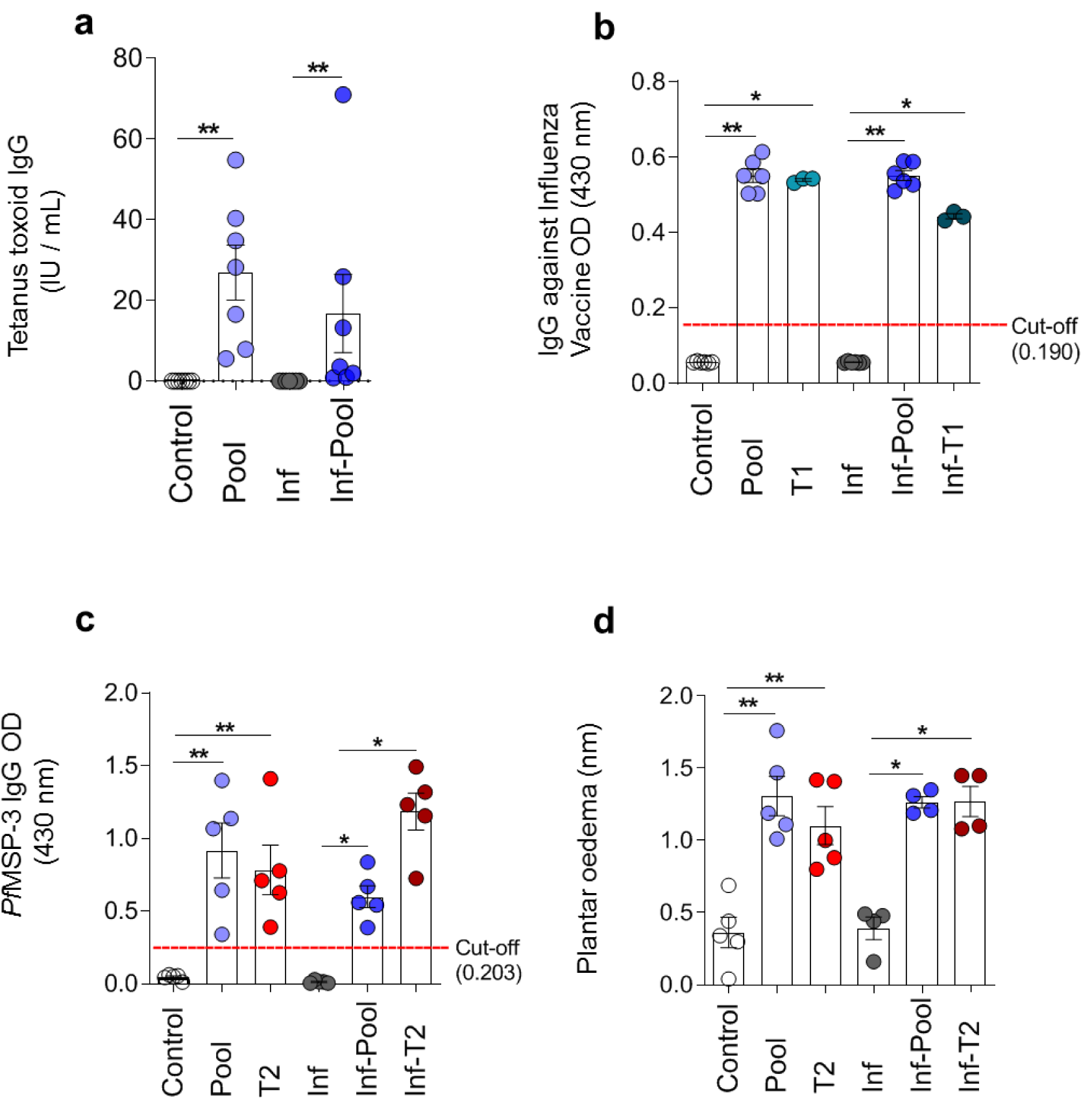
Immune stimulation with dT and influenza vaccines, *Pf*MSP-3 and OVA proteins triggers specific immune responses. Healthy or *Pb*A-infected (and treated) mice were immune stimulated, or not, with the strategies: Pool, T1 or T2. After behavioural evaluation, mice were randomly chosen for the analysis of the effectiveness of immune stimulation. Serum levels of (**a**) dT-specific IgG (n = 7), (**b**) Influenza-specific IgG (n = 3 - 6), and (**c**) *Pf*MSP-3-specific IgG (n = 5) were measured. **e**, Reaction to OVA was elicited by intradermal injection of the antigen in the footpad of the OVA-sensitized mice. Oedema was determined by measuring the thickness of the paw before and after inoculation (n = 4 - 5). Experimental groups: Control (non-infected / non-immune stimulated); Pool (non-infected / Pool-immune stimulated); T1 (non-infected / T1-immune stimulated); T2 (non-infected / T2-immune stimulated); Inf (infected / non-immune stimulated); Inf-Pool (infected / Pool-immune stimulated); Inf-T1 (infected / T1-immune stimulated); Inf-T2 (infected / T2-immune stimulated). Data are expressed as mean and s.e.m. \*\**P* < 0.01; \**P* < 0.05; Unpaired t-test with Mann-Whitney test was used.

**Extended data, Fig. 4.**
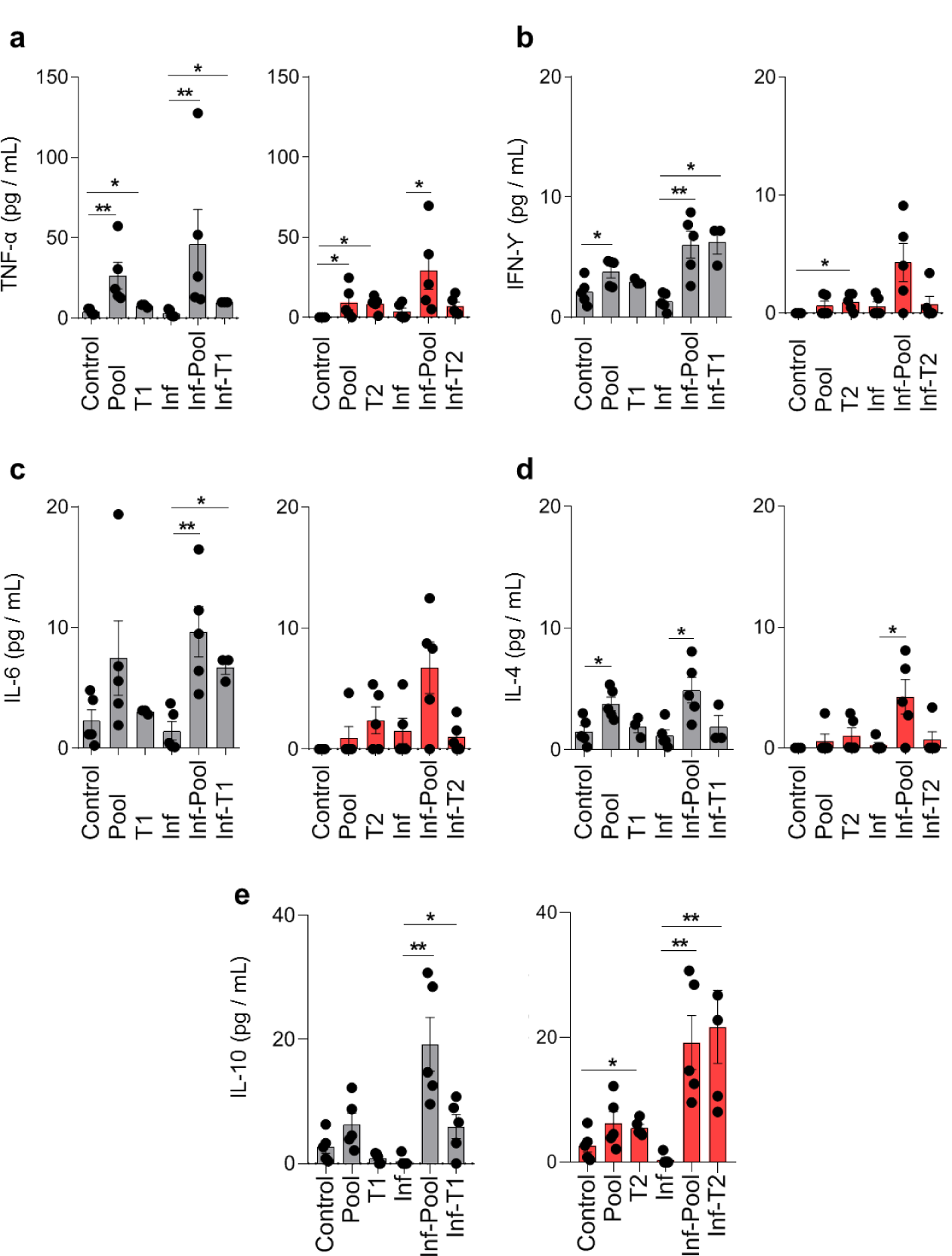
Immune stimulation with the different strategies elicits cellular responses measured by increased serum cytokine levels. Healthy or infected (and treated) mice were immune stimulated, or not, with the strategies: Pool, T1 or T2. Serum samples were collected after the behavioural evaluation (84 days after the end of CQ treatment), and levels of the cytokines (**a**) TNFα, (**b**) IFNγ, (**c**) IL-6, (**d**) IL-4 and (**e**) IL-10 were quantified by flow cytometry using cytometric bead array. Experimental groups: Control (non-infected / non-immune stimulated, n = 3 - 5); Pool (non-infected / Pool-immune stimulated, n = 5); T1 (non-infected / T1-immune stimulated, n = 3); T2 (non-infected / T2-immune stimulated, n = 5); Inf (infected / non-immune stimulated, n = 5); Inf-Pool (infected / Pool-immune stimulated, n = 5); Inf-T1 (infected / T1-immune stimulated, n = 3); Inf-T2 (infected / T2-immune stimulated, n = 5). Data are representative of three (Control, Pool, Inf and Inf-Pool groups) and one (T1, T2, Inf-T1 and Inf-T2 groups) independent experiments. Data are expressed as mean and s.e.m. \*\**P* < 0.01; \**P* < 0.05; Unpaired t-test with Mann-Whitney test was used.

**Extended data, Fig. 5.**
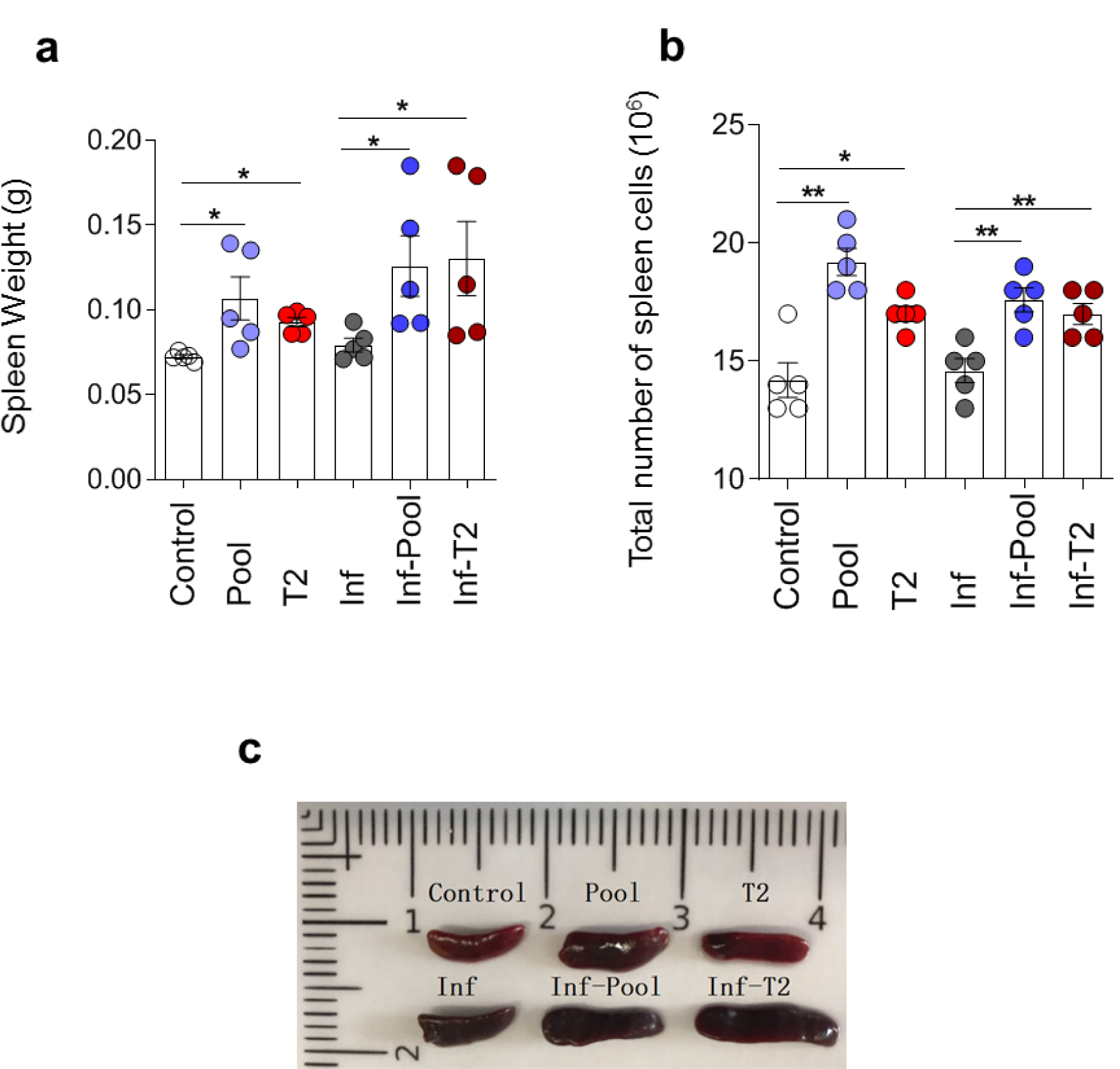
Splenic enlargement is observed after immune stimulation. Healthy or infected (and treated) mice were immune stimulated, or not, with the Pool or T2 strategy. Spleen weight (**a**) and total number of splenocytes (**b**) were evaluated at the end of the cognitive behavioural tasks (n = 5) (84 days after the end of CQ treatment). **c**, Representative photograph of Control, Pool, T2, Inf, Inf-Pool and Inf-T2 groups. Groups of infected mice showed a dark colour attributed to hemozoin, even more than two and a half months after infection. Experimental groups: Control (non-infected / non-immune stimulated); Pool (non-infected / Pool-immune stimulated); T2 (non-infected / T2-immune stimulated); Inf (infected / non-immune stimulated); Inf-Pool (infected / Pool-immune stimulated); Inf-T2 (infected / T2-immune stimulated). Data are mean and s.e.m. \*\**P* < 0.01; \**P* < 0.05; Unpaired t-test with Mann-Whitney test was used.

**Extended data, Fig. 6.**
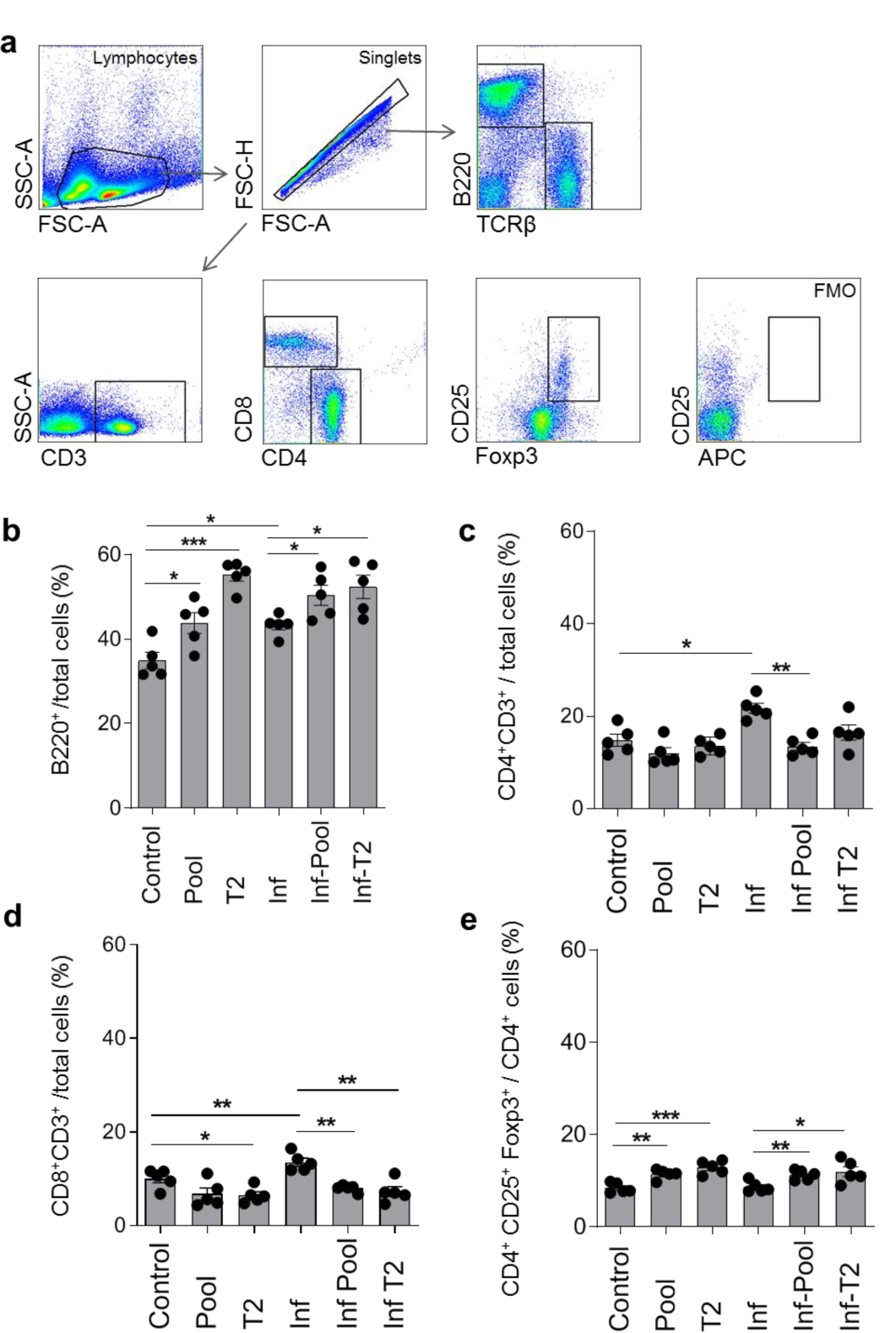
Stimulation of the immune system by the Pool and T2 strategies induces differentiation of Treg cells among the CD4 T cell population. Healthy or infected (and treated) mice were immune stimulated, or not, with the Pool or T2 strategy. Splenic lymphocytes subpopulations were analysed at the end of the cognitive behavioural tasks (84 days after the end of CQ treatment), when five mice were randomly chosen per group. **a**, Representative gating strategy to identify the populations of B cells (B220^+^), CD4 T cells (CD3^+^CD4^+^), CD8 T cells (CD3^+^CD8^+^) and Treg cells (CD3^+^CD4^+^CD25^+^Foxp3^+^) by flow cytometry. Percentage of B cells (**b**), CD4 T cells (**c**) and CD8 T cells (**d**) per spleen. **e**, Percentage of Treg cells among the CD4 T cells population. Experimental groups: Control (non-infected / non-immune stimulated); Pool (non-infected / Pool-immune stimulated); T2 (non-infected / T2-immune stimulated); Inf (infected / non-immune stimulated); Inf-Pool (infected / Pool-immune stimulated); Inf-T2 (infected / T2-immune stimulated). Data are expressed as mean and s.e.m. \*\**P* < 0.01; \**P* < 0.05; Unpaired t-test with Mann-Whitney test was used.

**Extended data, Fig. 7.**
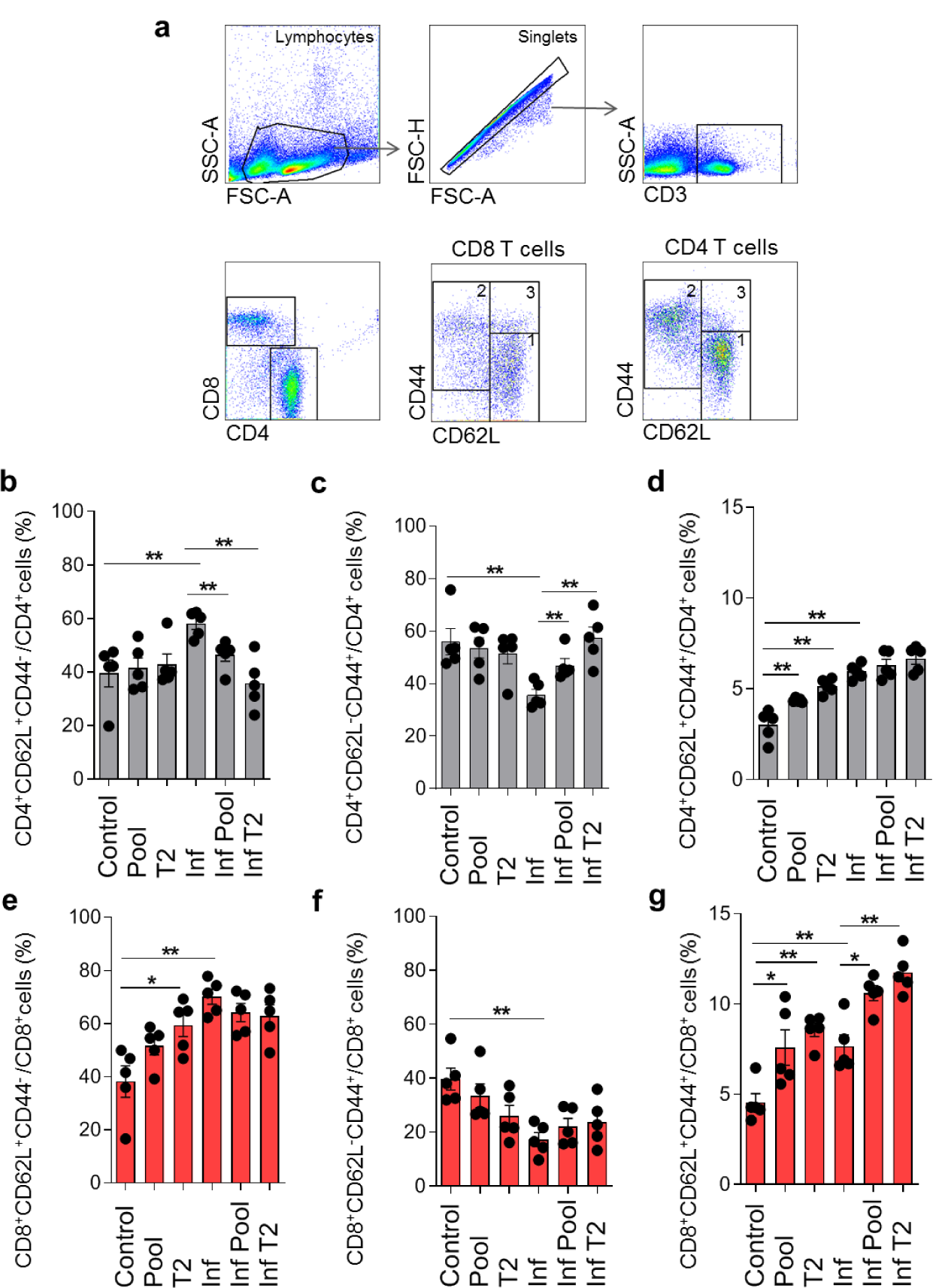
Effect of immune stimulation with the Pool and T2 strategies on the activation and memory phenotypes of CD4 and CD8 T cells. Healthy or infected (and treated) mice were immune stimulated, or not, with Pool or T2 strategy. Splenic lymphocyte subpopulations were analysed at the end of the cognitive behavioural tasks (84 days after the end of CQ treatment), when five mice were randomly chosen per group. **a**, Representative gating strategy to identify the subpopulations of naïve (gate: 1; CD44^-^CD62L^+^); effector / effector memory (gate: 2; CD44^+^CD62L^-^) and central memory (gate: 3; CD44^+^CD62L^+^) CD4 and CD8 T cells by flow cytometry. Percentage of naïve, effector / effector memory and central memory CD4 T cells (**b**-**d**) and CD8 T cells (**e**-**g**). Experimental groups: Control (non-infected / non-immune stimulated); Pool (non-infected / Pool-immune stimulated); T2 (non-infected / T2-immune stimulated); Inf (infected / non-immune stimulated); Inf-Pool (infected / Pool-immune stimulated); Inf-T2 (infected / T2-immune stimulated). Data are mean and s.e.m. \*\**P* < 0.01; \**P* < 0.05; Unpaired t-test with Mann-Whitney test was used.

